# Amyloid-β seeding effects are dependent on the presence of knock-in genes in the App^NL-G-F^ mice

**DOI:** 10.1101/2022.05.28.492382

**Authors:** Sean G. Lacoursiere, Jiri Safar, David Westaway, Majid H. Mohajerani, Robert J. Sutherland

## Abstract

Alzheimer’s disease (AD) is characterized by the prion-like propagation of amyloid-β (Aβ). However, the role of Aβ in cognitive impairment is still unclear. To determine the causal role of Aβ in AD, we intracerebrally seeded the entorhinal cortex of two-month-old *App*^*NL-G-F*^ mouse model with an Aβ peptide derived from patients who died from rapidly progressing AD. When the mice were three and six months of age, or one- and four-months following seeding, respectively, spatial learning and memory were tested using the Morris water task. Immunohistochemical labeling showed seeding with the Aβ seed increased plaque size one month following seeding, but reduced plaque counts four months following injection compared to the control seeded mice. A significant increase in microgliosis was found. However, we found no correlation between pathology and spatial performance. The results of the present study show that seeding human tissue with or without Aβ alters learning and memory ability, Aβ plaque deposition, plaque size, and microgliosis in the *App*^*NL-G-F*^ knock-in model, and these effects are dependent on the presence of a humanized *App* gene and the presence of Aβ in the seed. But these pathological changes were not initially causal in memory impairment.

## INTRODUCTION

The cause of AD may be due to the prion-like spread of Aβ which results in neuroinflammation, plaque deposition, and hyperphosphorylation of tau, ultimately causing synapse loss, brain atrophy (Bloom, 2014; Harper & Lansbury, 1997; Walker et al., 2018). The templated misfolding of native Aβ into misfolded Aβ provides the basis for experimental paradigms, providing an explanation for stereotypical patterns of deposition ascertained from “staging” human pathological material (Braak & Del Tredici, 2014; Eisele, 2013). Seeding Aβ is one method to cause reliable spread and deposition of Aβ and has become a useful tool to study the spread and aggregation of Aβ in mouse models (Friesen & Meyer-Luehmann, 2019; McAllister et al., 2020). The seeding properties of Aβ have been described independently by many labs, and studies have shown that Aβ comes in many varieties which differ in their aggregation and conformation properties and that Aβ from derived biological sources differs from synthetic peptides (Hatami et al., 2017; Jucker & Walker, 2018; Vandersteen et al., 2012; Ye et al., 2016).

Yet, there exists a paucity of studies examining the effects of Aβ seeding and deposition on cognitive decline in second generation mouse models – models that do not over express amyloid precursor protein. Previously our lab characterized the biochemical and behavioural changes that occur over time in the App^NL-G-F^ knock in mouse to address questions related to the development of characterized Aβ pathology in second generation mouse models (Mehla et al., 2019). Aβ aggregation, astrocytosis, a reduction in cholinergic and dopaminergic tone, and significant cognitive impairment all occurred at advanced ages. This model is appropriate as it develops AD-like pathology without over expressing APP, and is a proper host to study the effects of seeding Aβ and its prion-like propagation (Meyer-Leuhmann et al., 2006; Purro et al., 2018; Ruiz-Riquelme et al., 2018). Introducing Aβ and accelerating its deposition provides a window to study the effects human Aβ on learning and memory from aging and other pathological changes that naturally occur in this model.

The objective of the present experiments was to assess the behavioural and pathological effects of seeding human Aβ into the knock-in *App*^*NL-G-F*^ mouse. We intracerebrally injected a unique rapidly progressing Alzheimer’s disease Aβ seed (rpAD) isolated from human hippocampal tissue into the *App*^NL-G-F^ mice and their derivatives to determine the effects of Aβ deposition and microgliosis and spatial learning and memory. In this study we had three aims: 1) determine if seeding human hippocampal tissue with or without rpAD would impair spatial learning and memory in the *App*^NL-G-F^ mouse line; 2) if the two seeds would increase Aβ pathology elsewhere in the brain; 3) determine whether the Aβ aggregation correlated with cognitive impairments; 4) and to determine if this seeding exacerbated microgliosis.

## METHODS

Thirty homozygous negative *App*^NL-G-F^ mice (i.e., mice with an endogenous APP locus not perturbed by gene targeting; *App*^-/-^), ten heterozygotes *App*^NL-G-F^ (*App*^+/-^) and thirteen homozygous *App*^NL-G-F^ mice (*App*^+/+^), were used in this study (Table 1), with groups consisting of both male and females.

**Table 1.1.**
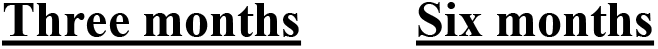
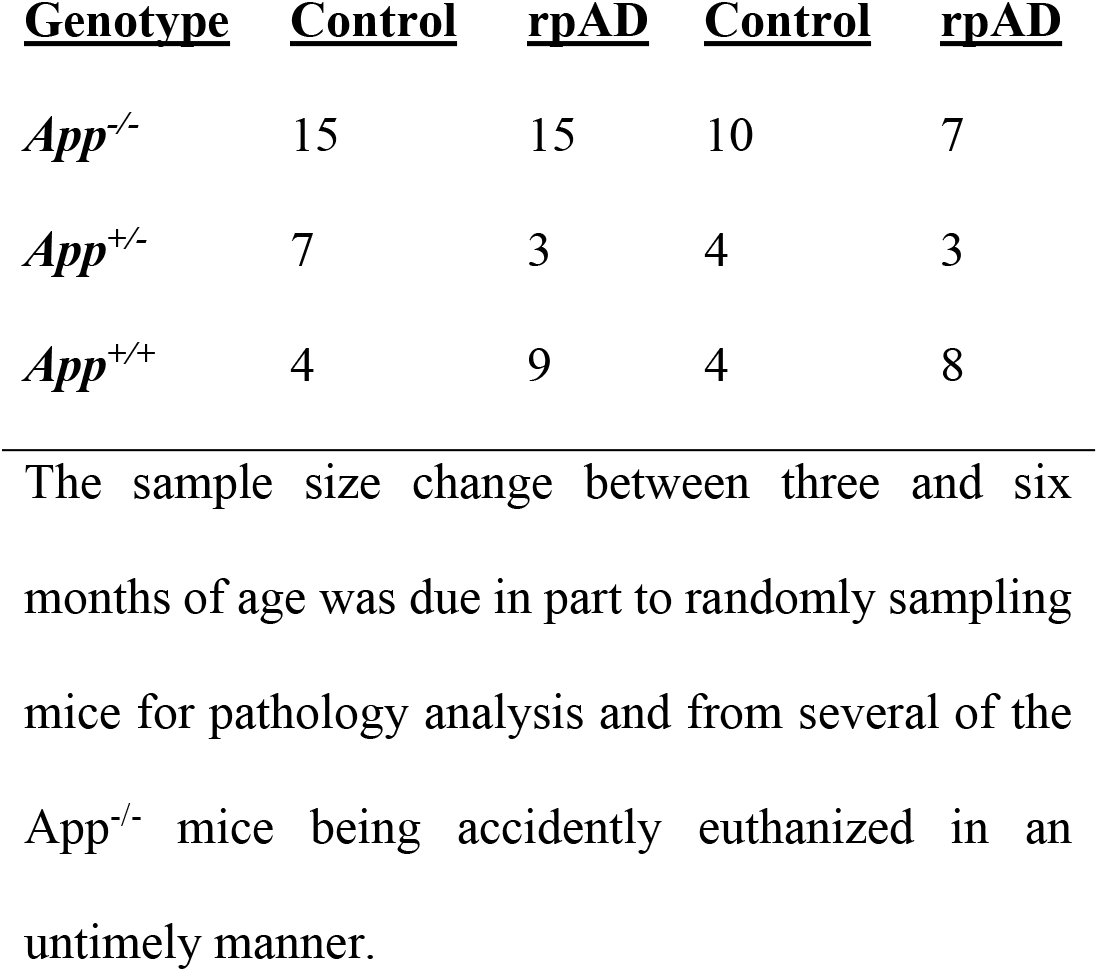
Final grouping of seed and genotype.

### Procedure and Statistics

The mice were pretrained on the MWT prior to surgery. At two months of age, the mice were intracerebrally seeded with a control or rpAD seed. One- and four-months following seeding or when the mice were three and six months of age, respectively, the MWT was used to test spatial memory. Following testing, the mice were randomly selected for perfusion and their pathology was assessed using immunohistochemistry. The behavioural data with repeated measures were analyzed using a two-way repeated measures ANOVA with experimental group as the independent measure and the training days as the repeated factor. A mixed effect 3-way ANOVA was used to analyze pathology data with Aβ plaque count and size, and activated microglia count as the dependent measures and seed, genotype, and age as independent measures. α was set to 0.05. Data presented as mean ± SEM unless otherwise stated.

## RESULTS

### Genotype and seed had unique effects on learning, memory, and Aβ plaque deposition and microgliosis in the App^NL-G-F^ mice

The *App*^-/-^, regardless of seed, developed no pathology (Fig 1.1 and Fig S1)) and were the only group that showed evidence of learning the MWT at both three and six months (Fig 1.2A and 1.3A). However, it was only the *App*^-/-^ C mice that showed evidence of learning and memory in the MWT at six months; the *App*^-/-^ rpAD seeded mice showed evidence of learning the task but did not show memory of the target location at six months (Fig 1.3A).

**Fig 1.1.**
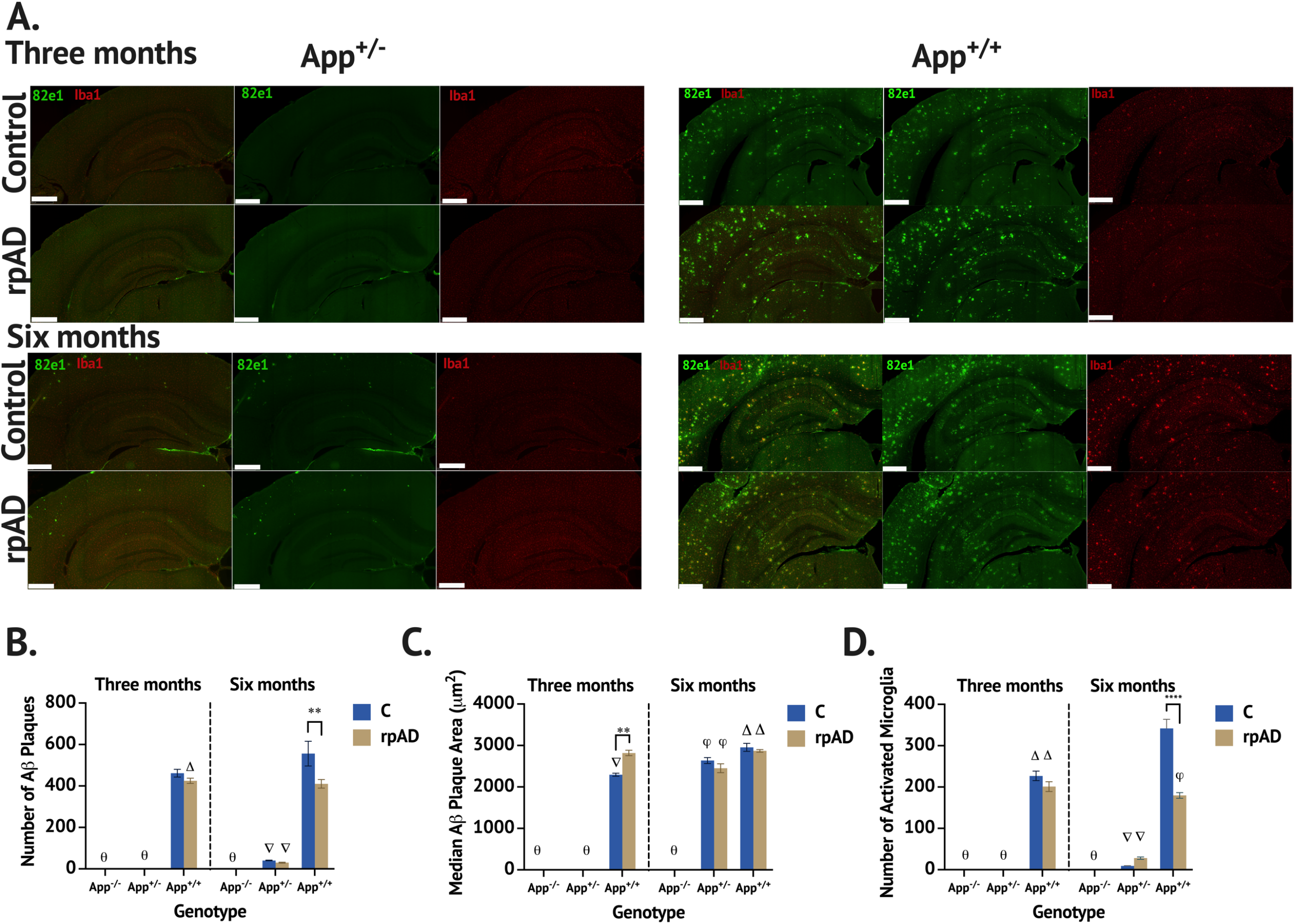
Intracerebral seeding altered plaque deposition and microgliosis in the brain of the *App*^*NL-G-F*^ mice that were unique to the seed, genotype, and age of the mouse. **A**. Photomicrographs of the hippocampus and cortex of the *App*^*+/-*^ and *App*^*+/+*^ mice at three (top panel) and six (bottom panel) months stained for Aβ (82e1) and activated microglia (Iba1). Scale bar set to 500 µm. **B**. Aβ plaque count following seeding was highly dependent on the genotype of the mouse and seed. Δ = significant difference between the *App*^*+/+*^ rpAD at three months and *App*^*+/+*^ control at six months *(p* = 0.0085). θ = significant difference from the *App*^*+/+*^ at three and six months (*p* < 0.0001). ∇= significant difference between the *App*^*+/-*^ C and rpAD seeded mice and the *App*^*+/+*^ C and rpAD seeded mice (*p* < 0.0001). **C**. Aβ plaque size was largest in the *App*^*+/+*^ mice compared to the *App*^+/-^ mice. The *App*^+/+^ rpAD seeded mice had significantly larger plaque at three months compared to the *App*^+/+^ C mice. θ = the *App*^*-/-*^ and *App*^*+/-*^ at three months had significantly smaller plaque compared to *App*^*+/-*^ at six months and *App*^*+/+*^ at three and six months. Δ = the *App*^*+/+*^ C and rpAD seeded mice had significantly larger plaque size compared to the *App*^*+/-*^ C and rpAD (*p* < 0.05). As shown, the *App*^*+/+*^ rpAD seeded mice had a significantly larger plaque compared to the *App*^*+/+*^ C seeded mice at three months. ϕ = significant increase in plaque size in the *App*^*+/-*^ C and rpAD seeded mice from three to six months (*p* < 0.0001) **D**. Activated microglia was significantly affected by genotype, seed, and age. θ = the *App*^*+/+*^ C and rpAD seeded mice had significantly more activated microglia at three and six months compared to the *App*^*-/-*^ at three and six months and the *App*^*+/-*^ mice at three months (*p* < 0.0001). Φ = significant difference between *App*^*+/+*^ C seeded mice at three months and *App*^*+/+*^ rpAD seeded mice at six months (*p* = 0.0025). ∇= significantly less activated microglia in the *App*^*+/-*^ C and rpAD seeded mice compared to *App*^*+/+*^ C and rpAD seeded mice at six months (*p* < 0.0001). Δ = significantly less activated microglia in the *App*^+/+^C and rpAD seeded mice at three months compared to the *App*^+/+^ C seeded mice at six months.

**Fig 1.2.**
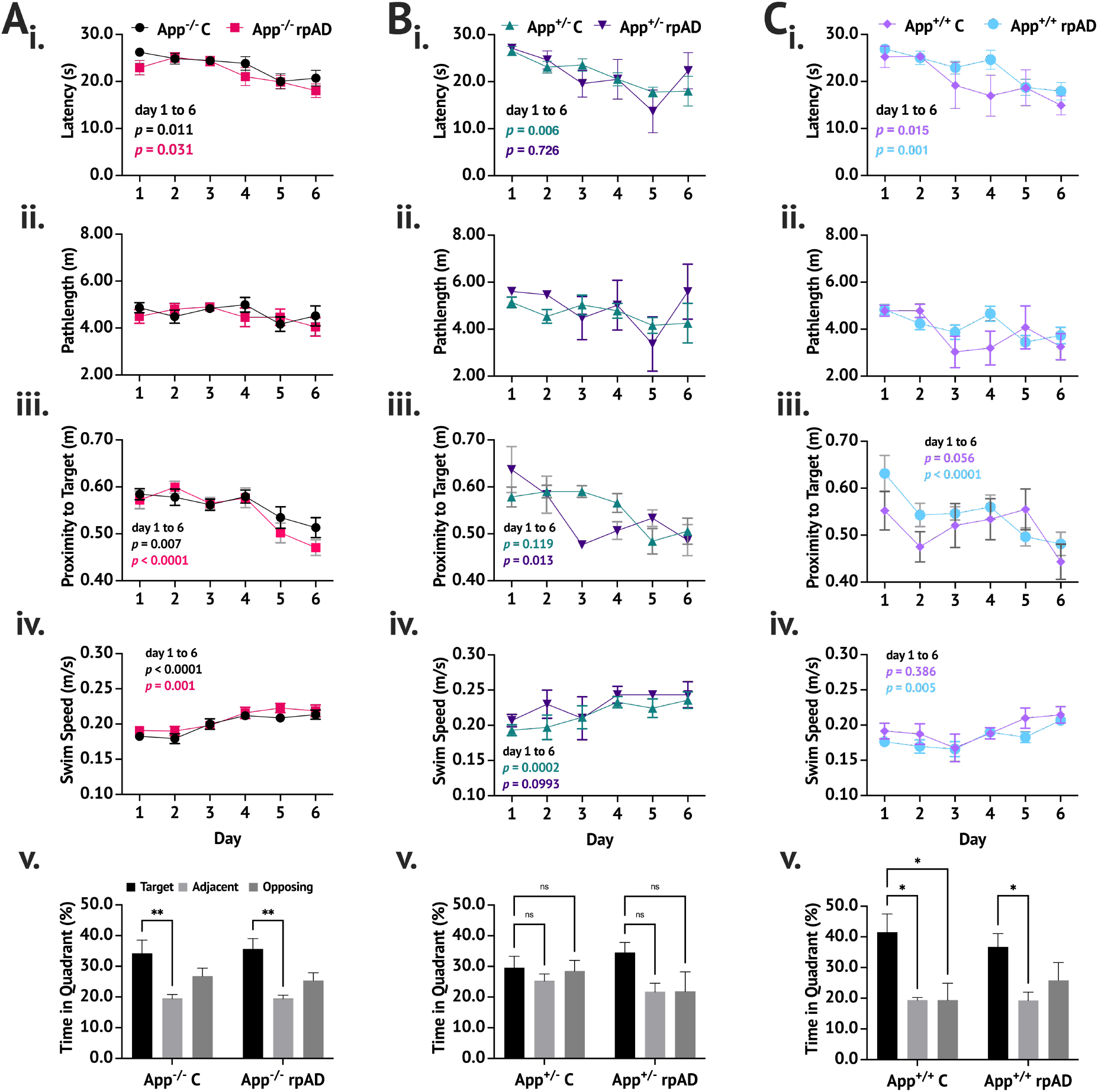
MWT performance at three months of age revealed no significant effect of seed but the *App*^+/-^ did not learn the task. **A**. The *App*^-/-^ C (n = 15) and *rpAD* (n = 15) seeded mice showed learning and memory of the target platform at three months. The *App*^-/-^ C and rpAD seeded mice reduced their latency (**i**.) and proximity to the target (**iii**.) but their pathlength did not change (**ii**.). The *App*^-/-^ C and rpAD mice showed significant preference for the target quadrant in the no-platform probe trial (**iv**.). **B**. The *App*^+/-^ C (n = 7) and rpAD (n = 4) seeded mice showed evidence of learning the task but did not show memory for the target. A consistent increase in swim speed was found in the *App*^-/-^ C, *App*^-/-^ rpAD, *App*^+/-^ C, and *App*^+/+^ rpAD seeded mice; the *App*^+/-^ C and rpAD seeded mice were significantly faster than the *App*^+/+^ rpAD mice (**iv**.). **C**. The *App*^+/+^ C (n = 4) and rpAD (n = 9) mice showed learning and memory at three months, reducing their latency to escape (**i**.) and proximity to the target (**iii**.) over training. The *App*^+/+^ C and rpAD mice spent significantly more time in the target quadrant compared to either the adjacent or opposing quadrants (**v**.). Data presented as mean ± SEM.

**Fig 1.3.**
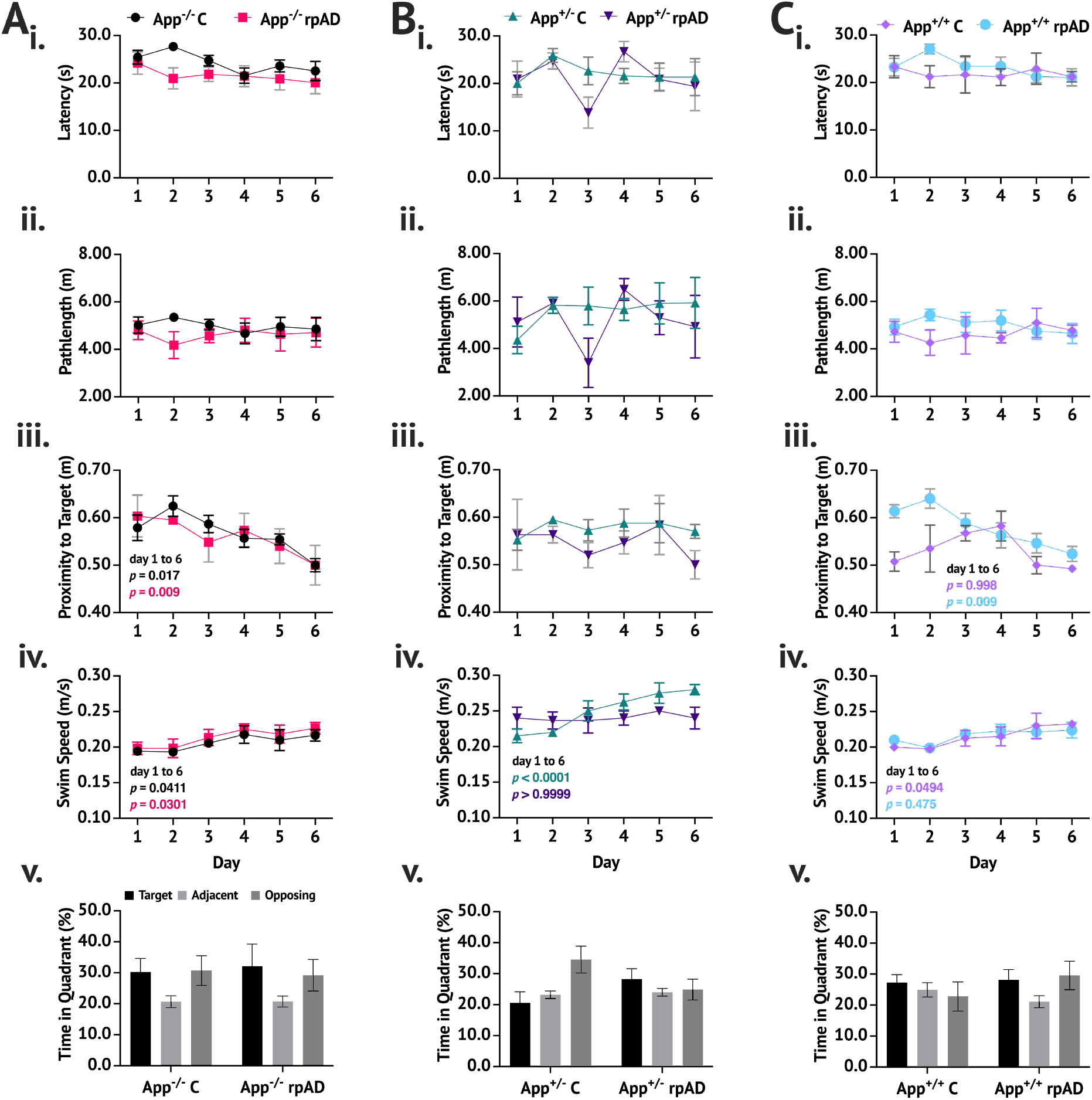
MWT performance at six months of age revealed no significant effect of seed but the App^+/-^ did not learn the task. The *App*^-/-^ C (n = 10) *App*^*-/-*^ rpAD (n= 7) and *App*^+/+^ rpAD (n = 8) seeded mice significantly reduced their proximity to the target over training (*p* < 0.05). The *App*^+/-^ C (n = 4), *App*^+/-^ rpAD (n = 3) and *App*^+/+^ C (n = 4) seeded mice showed no evidence of learning the task. **A**. MWT performance of *App*^-/-^ mice, **B**. The *App*^+/-^ mice showed no evidence of learning or memory at six months of age. **C**. The *App*^+/+^ mice showed a reduction in proximity to the target quadrant (**iii**.) but no reduction in latency (**i**.) or pathlength (**ii**.); they also showed no preference for the target quadrant in the no-platform probe trial (**v**.). Data presented as mean ± SEM.

**Fig 1.4.**
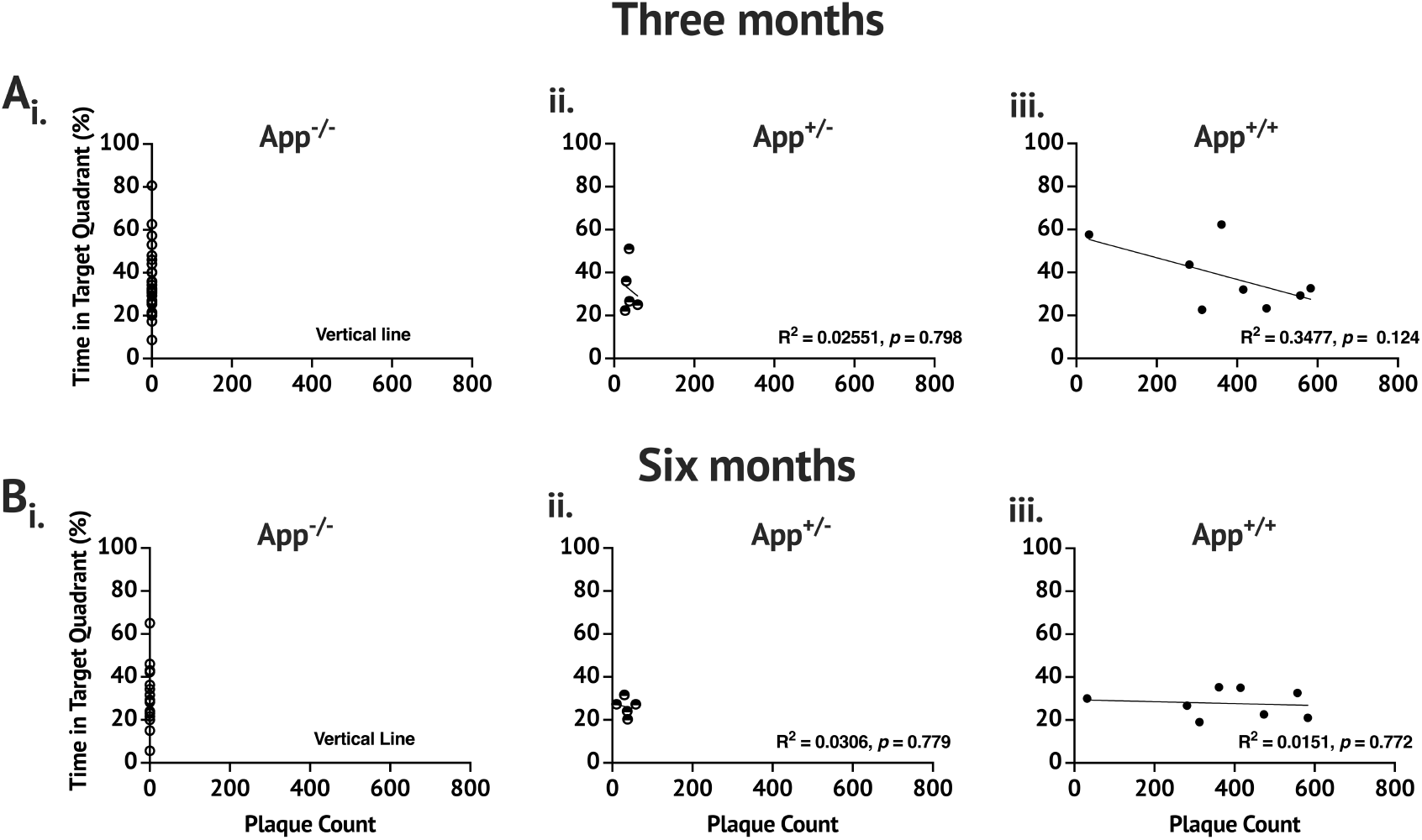
Correlation of Aβ plaque count and MWM no-platform probe trial performance. **A**. Three-month correlations of (**i**.) *App*^-/-^ mice; (**ii**.) *App*^+/-^ mice; (**iii**.) *App*^+/+^. **B**. Six-month correlations of (**i**.) *App*^-/-^ mice; (**ii**.) *App*^+/-^ mice; (**iii**.) *App*^+/+^. No significant correlations were found for any genotype.

The *App*^+/-^ C and rpAD mice did not develop plaque until six months of age (Fig 1B) and compared to the *App*^+/+^ C and rpAD mice, developed significantly fewer and smaller plaques, and developed significantly fewer activated microglia. However, the *App*^+/-^ C and rpAD mice showed a positive correlation between Aβ plaque and activated microglia (Fig 4C.). Despite developing significantly less pathology than the *App*^+/+^ C and rpAD mice, the *App*^+/-^ mice showed the poorest spatial learning and memory ability. The *App*^+/-^ C and rpAD mice did not show memory for the target location at three months of age (Fig 2.) and these mice showed no evidence of learning or memory of at six months of age (Fig 3.).

The *App*^+/+^ C and rpAD mice showed the greatest changes in plaque load and microgliosis following seeding (Fig. 1.1). At six months of age, the *App*^+/+^ C mice had a significantly greater plaque load and more activated microglia compared to the *App*^+/+^ rpAD seeded mice but the plaque size in the *App*^+/+^ rpAD seeded mice grew significantly quicker than the *App*^+/+^ C seeded mice. The *App*^+/+^ C seeded mice had more activated microglia at three and six months compared to the rpAD seeded mice at six months. At three months of age, the *App*^+/+^ C seeded mice did not show a positive correlation between Aβ plaque and activated microglia but the rpAD seeded mice did. Despite showing significant pathology at three months of age, the *App*^+/+^ C and rpAD seeded mice showed evidence of learning and memory (Fig. 1.2C). At six months however, the *App*^+/+^ rpAD seeded mice but not the *App*^+/+^ C seeded mice showed evidence of learning but neither *App*^+/+^ C and rpAD showed memory of the target location (Fig 1.3C).

### MWT one month post seeding

When the mice were three months of age, the *App*^-/-^ and *App*^+/+^ mice, regardless of seed, showed evidence of learning the location of the hidden platform in the MWT but the *App*^+/-^ mice did not (Fig 2). The mice significantly reduced their latency to escape [*F*(5, 235) = 15.19, *p* < 0.0001], pathlength swam [*F*(5, 235) = 3.845, *p* = 0.0023] and average proximity to target [*F*(5, 235) = 14.41, *p* < 0.0001] over training days. Aside from the *App*^*+/-*^ mice having a significantly faster swim speed compared to the *App*^*-/-*^ and *App*^*+/+*^ mice [*F*(2, 47) = 7.167, *p* = 0.0019], no group differences were found for latency [*F*(5, 47) = 1.190, *p* = 0.3285], pathlength [*F*(5, 47) = 2.306, *p* = 0.0592], or proximity [*F*(5, 47) = 0.9007, *p* = 0.4886]. However, a significant day x group interaction in proximity was found [*F*(25, 235) = 1.690, *p* = 0.0246]. The interaction is due to the day at which a significant reduction in proximity was first found in the group of mice. For example, the *App*^-/-^ C mice did not show a significant reduction until the sixth day of training (*p* = 0.006), whereas the *App*^+/+^ rpAD mice showed a significant reduction by the second day (*p* = 0.01)

In the no-platform probe trial, the mice spent significantly more time in the target quadrant compared to the non-target quadrants [*F*(2, 94) = 12.75, *p* < 0.0001]. No differences were found between groups [*F*(5, 47) = 0.3869, *p* = 0.8553]. The *App*^*+/-*^ control and rpAD seeded mice did not spend significantly more time in the target quadrant compared to the opposing or adjacent quadrants (*p* > 0.05) whereas the *App*^*-/-*^ and *App*^*+/+*^ mice all spent significantly more time in the target quadrant than in the adjacent quadrants. The *App*^*+/+*^ control seeded mice also spent significantly more time in the target quadrant compared to the opposing quadrant (Fig 1.1Cv.).

### MWT four months post seeding

At six months of age, only the *App*^-/-^ C mice showed evidence of learning and memory of the hidden platform. The mice showed a significant reduction in proximity to the target [*F*(5, 140) = 6.186, *p* < 0.0001]. The *App*^*-/-*^ C (*p* = 0.0152), *App*^*-/-*^ rpAD (*p* = 0.0081), and *App*^*+/+*^ rpAD (*p* = 0.0077) were the only mice that significantly reduced their proximity from the first to last day of training. The mice showed no reduction in latency [*F*(5, 140) = 2.060, *p* = 0.0740] or pathlength [*F*(5, 140) = 0.715, *p* = 0.6132]. Overall, the *App*^*+/-*^ mice had a significantly faster swim speed compared to the *App*^*-/-*^ and *App*^*+/+*^ mice [*F*(2, 28) = 8.834, *p* = 0.0011].

In the no-platform probe trial, the mice spent similar time in the target quadrant and non-target quadrants [*F*(2, 56) = 2.205, *p* = 0.1197], spending similar times between groups [*F*(5, 28) = 1.161, *p* = 0.3527]. One mouse in the *App*^*-/-*^ C group was found to spend the most time in the target quadrant at three months of age compared to all other mice, spending 80.6% of its time in the target quadrant. However, at six months of age, this mouse spent only 5.6% of its time in the newly trained target quadrant but spent 61.3% of time in the previously learned target quadrant (see Supplementary figures 2 – 7 for individual mouse swim paths during three- and six-month probes). If this mouse is removed from the analysis, it is found that the *App*^*-/-*^ control seeded mice spent significantly more time in the target quadrant than what chance would predict [*t*(7) = 2.362, *p* = 0.0251], however, compared to the adjacent and opposing quadrant the mice did not.

To assess whether the mice had remembered the previously learned target quadrant and the newly acquired quadrant, the time spent in the previously learned and recently learned target quadrants compared to the non-target quadrants were averaged. The mice spent significantly more time in the target quadrants than in the non-target quadrants [*F*(1, 28) = 11.26, *p* = 0.0023]. Despite no significant group differences in this measure [*F*(5, 28) = 0.4564, *p* = 0.8050] only the *App*^*-/-*^control seeded mice spent significantly more time in the new and old target quadrants compared to the adjacent non-target quadrants (*p* = 0.0212).

At six months the *App*^*+/-*^ mice were not able to learn the location of the hidden platform across training and did not remember its location in the no-platform probe trial. The control seed caused greater impairment in the *App*^*+/+*^ C seeded mice had greater impairment than the *App*^+/+^ rpAD seeded mice. The *App*^*-/-*^ C seeded mice were the only group to spend significantly more time in both previously learned and recently acquired target quadrants compared to the non-target quadrants suggesting they learned the target location and remembered the old target location. The *App*^-/-^ rpAD seeded mice showed impairment compared to the *App*^-/-^ C seeded mice.

Overall, the *App*^-/-^ and *App*^+/+^ mice were able to learn the task at three and six months of age and regardless of seed. The *App*^+/-^ mice showed impaired memory at three and six months – at three months these mice showed evidence of learning but no evidence of remembering the task; at six months, no evidence of learning or memory was found. Despite the group differences found at three months, neither the seed [*F*(1, 47) = 0.0141, *p* = 0.906] or genotype [*F*(2, 47) = 0.692, *p* = 0.506] was found to have a significant effect on time spent in the target quadrant. No effect of seed [*F*(1, 27) = 0.3730, *p* = 0.547] or genotype [*F*(2, 27) = 1.523, *p* = 0.236] was found at six months.

#### Aβ plaque deposition

Aβ plaque deposition throughout the brain was significantly affected by genotype [*F*(2, 520) = 562.7, *p* < 0.0001] and seed [*F*(1, 520) = 5.077, *p* = 0.025] and the genotype x seed interaction was significant [*F*(2, 520) = 5.323, *p* = 0.005]. Age, however, was not a significant factor on Aβ plaque count [*F*(1, 520) = 3.115, *p* = 0.078]. The *App*^*-/-*^ control and rpAD seeded mice never developed plaque. At three months, the *App*^*+/+*^ C and rpAD seeded mice had similar plaque load (461 ± 18.6 and 425.1 ± 12.44 plaques per section, respectively; *p* = 0.994) and a significantly greater plaque load than both the *App*^*-/-*^ and *App*^*+/-*^ mice (*p* < 0.0001). At six months of age, the *App*^*+/-*^ C and rpAD seeded mice began developing Aβ plaque, but this was significantly less than the *App*^*+/+*^ C and rpAD seeded mice at both three and six months.

The size of the individual plaques was significantly affected by the genotype [*F*(2, 456) = 487.8, *p* < 0.0001] and age [*F*(1, 184) = 314.8, *p* < 0.0001] of the mouse and a significant genotype x age interaction was founds [*F*(2, 184) = 202.4, *p* < 0.0001]. The seed itself did not have a significant effect on plaque size [*F*(1, 456) = 0.4972, *p* = 0.4811], but a significant age x seed interaction was found [*F*(1, 184) = 6.642, *p* = 0.0107]. At three months of age the *App*^*+/+*^ rpAD mice had significantly larger plaque compared to the App^+/+^ C seeded mice (*p* = 0.003) but at six months, no significant difference was found (*p* > 0.9999). The plaque size in the *App*^*+/+*^ rpAD seeded mice at six month were also significantly larger than the plaque in three-month-old *App*^+/+^ C seeded mice (*p* < 0.0001). The plaque grew significantly from three to six months in the *App*^*+/+*^control mice (*p* < 0.0001). The plaque size in the *App*^*+/+*^ rpAD seeded mice was not significantly different at three and six months (*p* = 0.825).

These results show that the rpAD seed increased the size of Aβ plaque quickly up to a maximal size in the *App*^*+/+*^ mice but over time, the plaque in the App^+/+^ C seeded brain grew to similar size. The plaque size in the *App*^*+/-*^ C and rpAD seeded mice grew from a point after three months to six months but was still significantly smaller compared to the *App*^*+/+*^ C and rpAD seeded mice. Seeding did not have as great of an effect on plaque count or size in the *App*^*+/-*^ mice. However, the results of the *App*^*+/-*^ seeded mice suggest that plaque can grow quickly to a roughly similar size or to a maximal size despite showing delayed onset.

#### Microglia

The activated microglia count was significantly affected by the genotype [*F*(2, 485) = 361.8, *p* < 0.0001], seed [*F*(1, 485) = 7.89, *p* = 0.005], and age [*F*(1, 158) = 6.425, *p* = 0.012]. Interactions between genotype x seed [*F*(2, 485) = 17.96, *p* = <0.0001], genotype x age [*F*(2, 158) = 3.670, *p* = 0.028], age x seed [*F*(1, 158) = 6.200, *p* = 0.014] and genotype x age x seed [*F*(2, 158) = 13.69, *p* < 0.0001] were found.

The *App*^*-/-*^ mice did not show activated microglia and the *App*^*+/-*^ mice showed activated microglia only at six months of age. Yet, compared to the *App*^*-/-*^ mice and three-month-old *App*^*+/-*^ mice, this increase was not significant, regardless of seed. At three months, the *App*^*+/+*^ control and rpAD seeded mice had similar counts of activated microglia with 226.95 ± 11.43 and 201.12 ± 11.80, respectively (*p* = 0.9982) but at six months, the *App*^*+/+*^ control seeded mice had significantly more activated microglia compared to the three- and six-month *App*^+/+^ rpAD seeded mice (*p* < 0.0001) and the three-month-old *App*^*+/+*^ control mice. The *App*^*+/+*^ control and rpAD seeded mice had significantly more activated microglia compared to the *App*^*-/-*^ and *App*^*+/-*^ mice at three and six months (Fig 1.1D.).

#### Aβ plaque deposition and no-platform probe trial performance were not corelated

A correlation of time in target quadrant during the no platform probe trial at three and six months with the number of Aβ plaque deposits in the *App*^*-/-*^, *App*^*+/-*^ and *App*^*+/+*^ control and rpAD seeded mice showed no significant correlation, suggesting that plaque deposition was not associated with performance (Fig 1.4.).

#### Aβ plaque burden and activated microglia are highly correlated

The Aβ plaque and activated microglia count for each brain section was correlated (Fig 1.5.). Aside from the exception of the three-month *App*^*+/+*^ control seeded mice, a strong positive correlation was found between activated microglia and Aβ plaque count in both the control and rpAD seeded *App*^*+/+*^ and *App*^*+/-*^ mice. Despite the *App*^*+/-*^ control and rpAD seeded mice having significantly less Aβ plaque and activated microglia, a significantly positive correlation between microglia and Aβ plaque deposition was found at six months (Fig. 1.5.).

**Figure 1.5.**
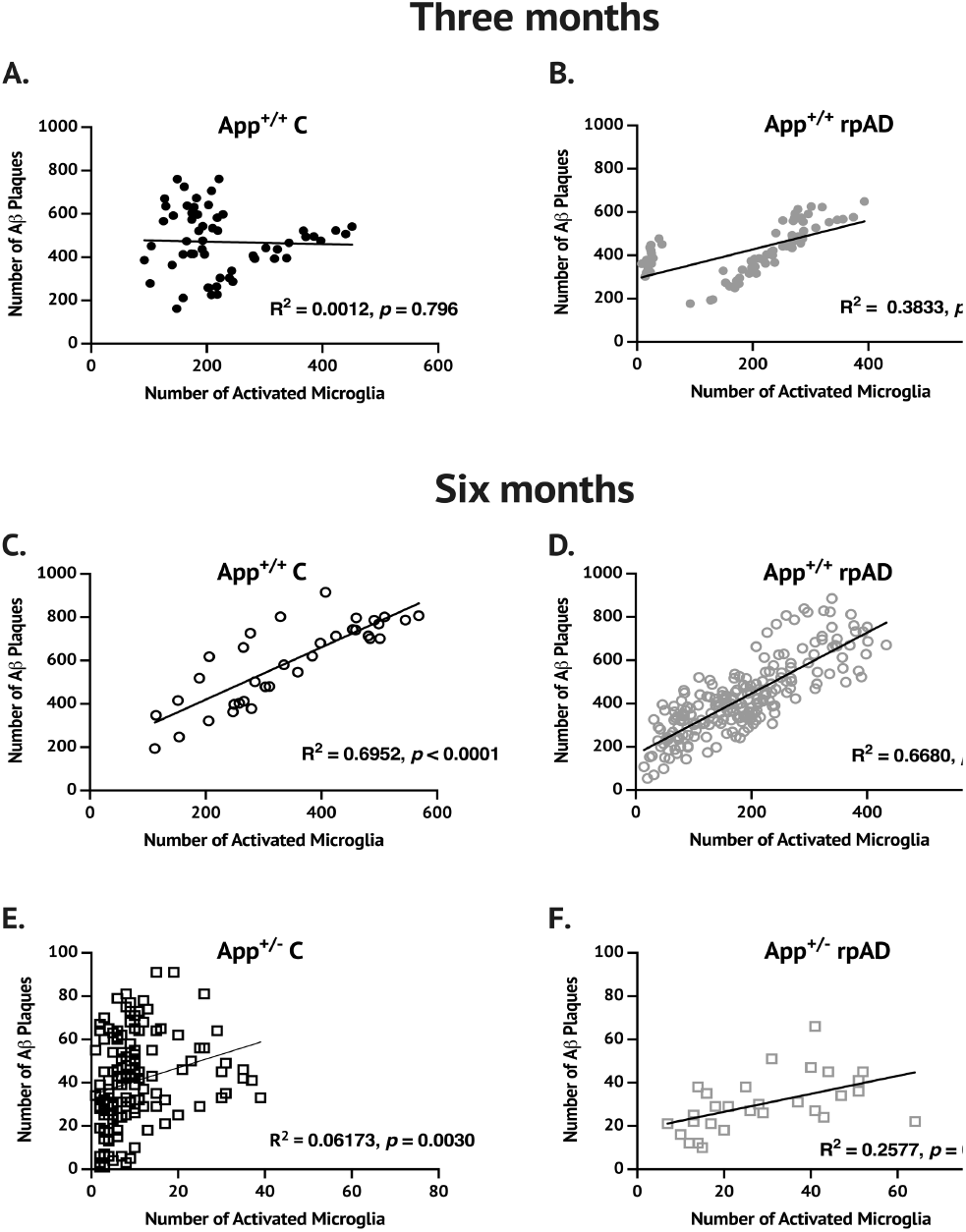
Correlation of Aβ plaque and activated microglia count in the *App*^*+/-*^ and *App*^*+/+*^. At each time point tested and for each seed and genotype except for the three-month *App*^+/+^ control seeded mice **(A)**, a significant positive correlation between Aβ plaque and activated microglia was found. The *App*^-/-^ mice had no plaque or microglia. Correlation was only done on mice with non-zero pathology values.

## DISCUSSION

The results of the present study show that seeding human tissue with or without Aβ alters learning and memory ability, Aβ plaque deposition, plaque size, and microgliosis in the *App*^NL-G-F^ knock-in model, and these effects are dependent on the presence of humanized *App* genes and the presence of Aβ in the seed.

### Aβ deposition and cognitive performance

Aβ deposition has been found in the *App*^*NL*-*G-F*^ mice at two – three and a half months (Latif-Hernandez et al., 2019; Saito et al., 2014), four months (Jafari et al., 2018) and after six months (Mehla et al., 2019). In these reports, the extent of Aβ aggregation is associated with cognitive deficits; such as in delay discounting, place preference, and attention control (Masuda et al., 2016), spatial learning and memory, fear conditioning, and object memory (Mehla et al., 2019), and levels of anxiety (Latif-Hernandez et al., 2019). Seeding Aβ has also been shown to impair performance in the MWT (Ziegler-Waldkirch et al., 2018). We predicted that cognitive decline would therefore correlate with Aβ pathology. However, the initial Aβ deposition and microgliosis found in our study did not correlate with learning and memory performance in the MWT. However, the only mice in our study that showed significant learning and memory were the *App*^-/-^ C, which had no plaque aggregation or seeded Aβ in their brain.

### Seeding in the *App*^*NL-G-F*^ mouse

Following seeding, both the control and rpAD seed increased Aβ plaque deposition with the rpAD seed causing plaques to grow larger sooner. Furthermore, we found an interesting dynamic between the size of plaque, the number of activated microglia, and the time following seeding in the App^NL-G-F^ mice. Despite these changes in pathology, no correlation was found between Aβ plaque deposition and microgliosis and learning and memory performance.

Accelerating Aβ plaque deposition in mouse models through Aβ seeding is a well characterized phenomenon (Burwinkel et al., 2018; Eisele et al., 2010; Rosen et al., 2012; Ye et al., 2016) (see (McAllister et al., 2020) for review). The seeding effect is due to templated misfolding of native Aβ species due to the conformational properties of the seed (Come et al., 1993; Eisele, 2013). We found that seeding Aβ accelerated the growth of plaque size in the brain of homozygous positive App^NL-G-F^ mice compared to control tissue seeded brains but the control tissue caused a greater increase in plaque number and activated microglia compared to Aβ seeded brains. Seeding Aβ may be attenuating creation of new plaque by promoting plaque growth and sequestration of Aβ (Liu et al., 2015) compared to control tissue seeded brains.

The *App*^-/-^ C or rpAD seeded mice did not develop plaque or microgliosis. Without the proper host, one lacking humanized Aβ expression, the seeding effect is not observed (Meyer-Leuhmann et al., 2006). The *App*^-/-^ C seeded mice were able to learn the MWT but the *App*^-/-^ rpAD mice were not, suggesting the rpAD is impairing memory despite showing no Aβ plaque deposition, soluble Aβ, or microgliosis following seeding of either homogenate (Fig S2). The *App*^+/-^ mice showed some Aβ plaque deposition and microgliosis but this was significantly attenuated compared to the *App*^+/+^ mice. Our results suggest that extent of amyloidosis from seeding is dependent on number of knocked-in mutations but that amyloidosis and microgliosis levels did not correlate with cognitive decline as both heterozygote and homozygous positive mice showed impairment at six months; or, any pathology or damage from Aβ can lead to cognitive decline.

### Spread of Aβ through microglia

Several mechanisms have been proposed describing how Aβ spreads through the brain. Neuron to neuron transport (Domert et al., 2014), proximity based diffusion (Mezias & Raj, 2017), or de novo amyloidosis (McAllister et al., 2020) have been suggested. One new line of thought is that the propagation and spread of Aβ in sporadic AD is thought to be initiated by the mechanisms underlying inflammation and infection (Ahmad et al., 2019; Heneka et al., 2018; Moir et al., 2018; Venegas et al., 2017). Microglia are the central nervous system resident immune cells, phagocytosing dead cells and fibrillar Aβ but uptaking other components through pinocytosis. It has been hypothesized that free, neuronally derived Aβ is internalized and aggregated in microglia and this aggregated Aβ is deposited into the extracellular space contributing to the initial formation of plaque (Spangenberg et al., 2019). Microglial elimination was found to impair plaque formation and the reemergence of microglia can seed and form plaques suggesting that plaque onset relies on the presence of microglia (Spangenberg et al., 2019). Microglia can rapidly take up and release both fibrillar and soluble Aβ, but with little degredation of the Aβ (Chung et al., 1999). Activated microglia surrounding Aβ plaques can take up Aβ but excessive uptake of Aβ can result in microglial death, resulting in the release of accumulated Aβ into the extracellular space and contributing to plaque growth (Baik et al., 2016).

Aβ, while categorized as a toxic protein, is an antimicrobial peptide (Soscia et al., 2010) shown to sequester toxic proteins of the same species (Liu et al., 2015), viruses (Eimer et al., 2018) and bacteria (Dominy et al., 2019). Microglia may use Aβ peptides to stop potential infection by actively creating plaque around damaged areas (Baik et al., 2016) and infectious agents (Eimer et al., 2018; Kumar et al., 2016); suggesting that the microglia may be using Aβ to increase the efficiency of the immune response in healthy brain, but in the mouse model, this may look like a systemic infection. Seeded Aβ may be internalized by microglia in an attempt to reconcile the increased Aβ levels as microglia try to remove Aβ. Primary microglia spontaneously move through microfluidic channels up to distances of 500 μm in approximately 12 hours, with an average speed of 0.66 μm/min (Amadio et al., 2013) and migrate towards Aβ plaque (Bolmont et al., 2008). Following the breakdown and uptake of Aβ plaque by microglia, microglia could migrate away from the original plaque location, and deposite Aβ into the extracellular space following, facilitating the spread of Aβ throughout the brain following microglia death (Baik et al., 2016; Bolmont et al., 2008; Chung et al., 1999). Due to the little degradation of Aβ that appears to be happening in microglia (Chung et al., 1999), an infectious Aβ oligomer could be taken up and spread from the plaque location.

## Limitations

There were several limitations to this study. First, the focus was on early disease effects on cognition and, specifically only on spatial navigation learning and memory. In only testing spatial navigation, we are unable to make conclusions on the effects of seeding, age, or genotype in different cognitive domains. However, the MWT was originally designed to test the function of the HPC in memory and one of the earliest impairments in AD is found in HPC memory. In future studies additional behavioural tests should be included or a novel home cage-based assessment of rodent’s behaviour (Contreras et al., 2022; Singh et al., 2019) may offer insights into phenotypical expression of mouse genotypes, aging, and seeding not found in traditional rodent behavioural testing. Second, we used repeated testing which has been avoided in previous studies (Mehla et al., 2019); but it has been argued by others that repeated testing is actually a better representative of clinical AD testing (Janus et al., 2000). Third, we did not focus on soluble Aβ and instead focused on Aβ plaque load and characteristics throughout the brain and how this was associated with microglia. We therefore cannot make any conclusions on how seeding affected soluble Aβ. Lastly, the loss of mice between the three- and six-month time points resulted in the completed study having less statistical power than initially designed, preventing us from finding small effects.

## Conclusion

In this study we seeded the *App*^*NL-G-F*^ knock-in mice with human tissue with or without human Aβ to determine the effects on Aβ plaque deposition, microgliosis, and learning and memory. We show that seeding is dependent on the presence of humanized *App* in a gene-dose response and on the presence of Aβ in the seed. Our results suggest that amyloidosis and microgliosis represent an immune response in brain and the response is dependent on the host and type of foreign agent introduced. The relationship between Aβ plaque deposition and microglial activation may provide alternative routes for therapy in treating AD. Future research will have to further examine the relationship between Aβ plaque deposition and microgliosis.

## SUPPLEMENTARY FIGURE

**Fig S1.1.**
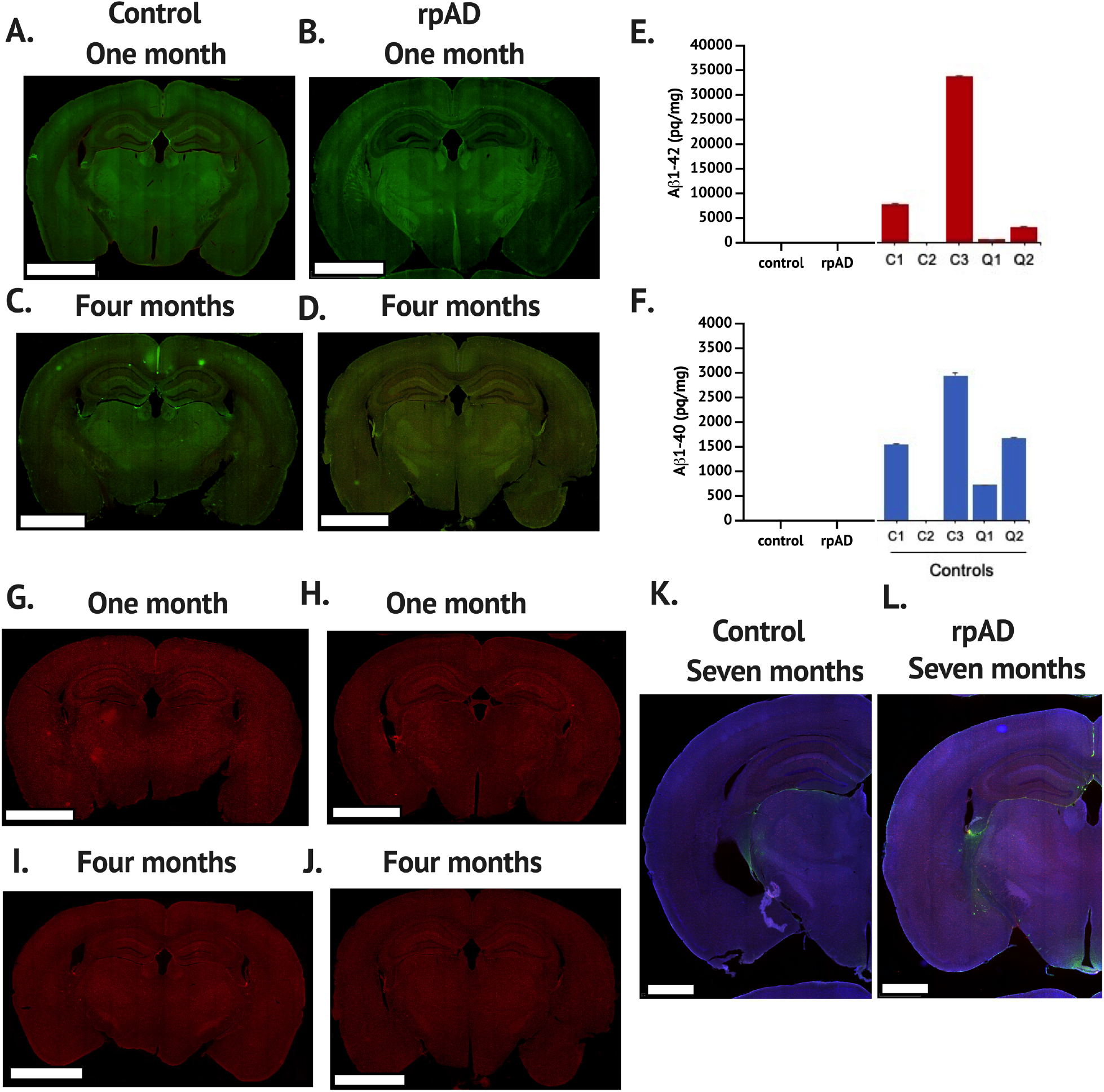
The *App*^-/-^ mice had no Aβ deposition, or microgliosis following seeding with either the C or rpAD seed at three or six months of age, or one and four months post seeding, respectively. No soluble Aβ_1-42_ or Aβ_1-40_ was found at six months of age in the *App*^-/-^ C or rpAD seeded mice, Furthermore, no pathology was found up to seven months following seeding or nine months of age.

**Fig S1.2.**
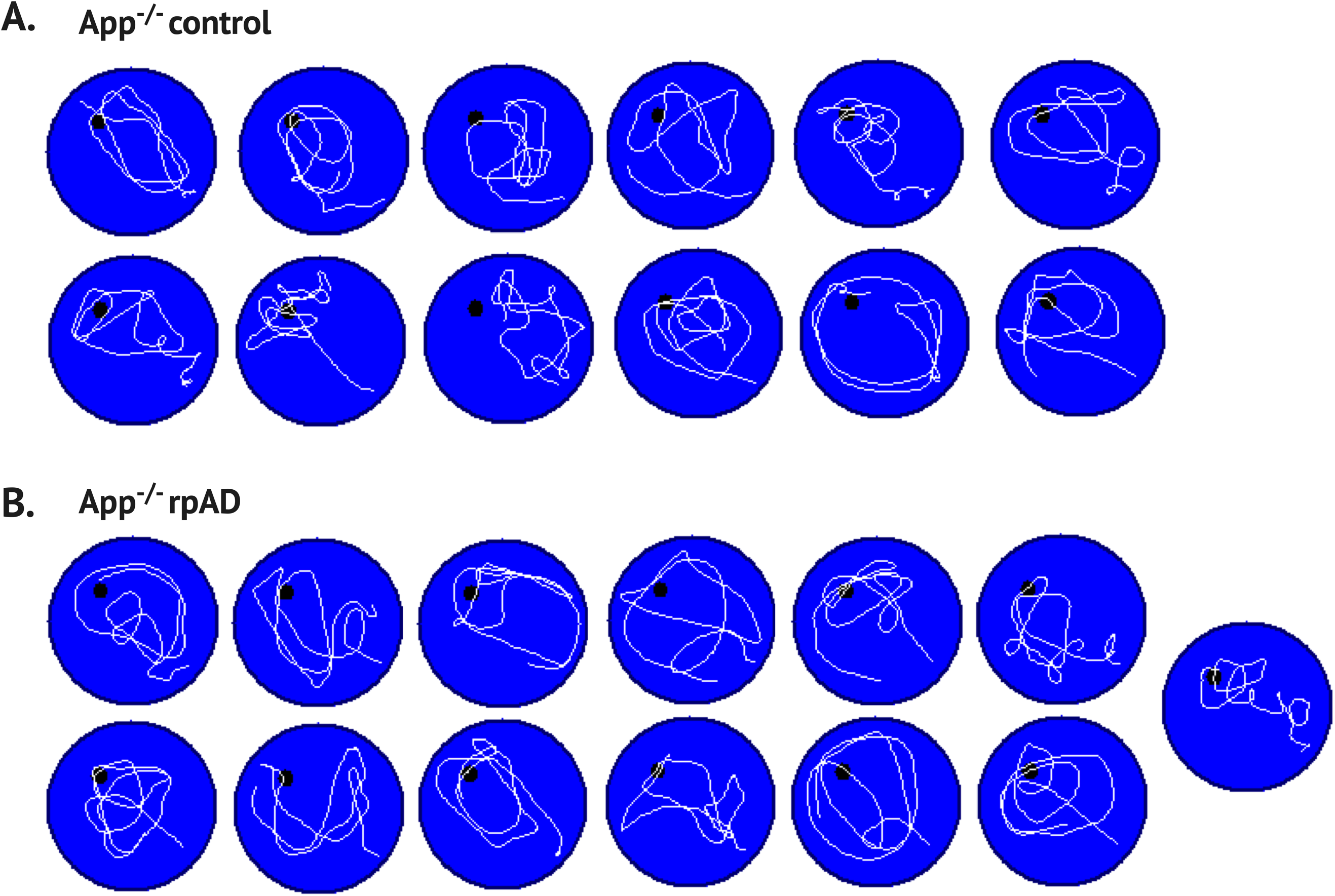
Swim paths for the App^-/-^ control (A) and rpAD (B) seeded mice at three months in the no-platform probe trial.

**Fig S1.3.**
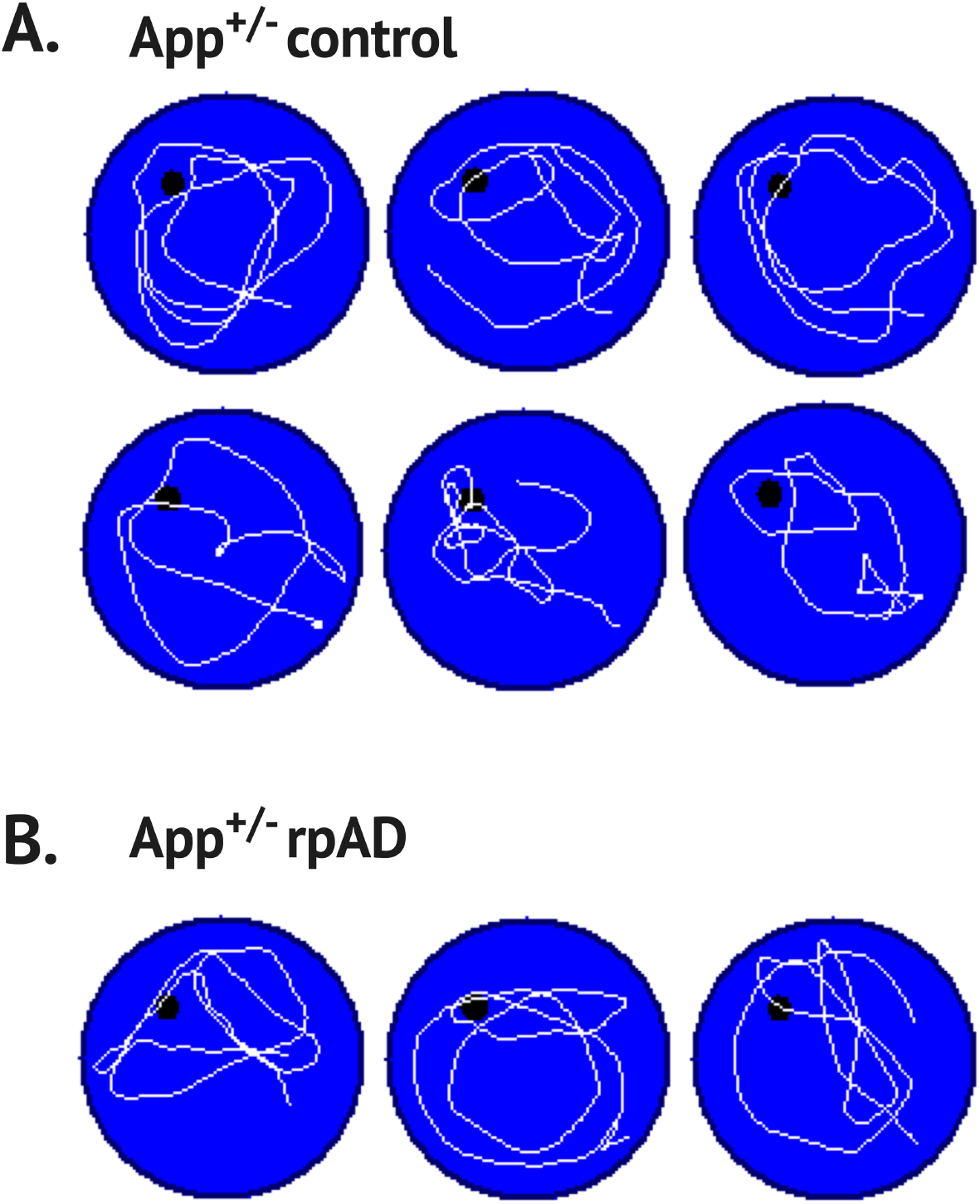
Swim paths for the App^+/-^ control (A) and rpAD (B) seeded mice at three months in the no-platform probe trial

**Fig S1.4.**
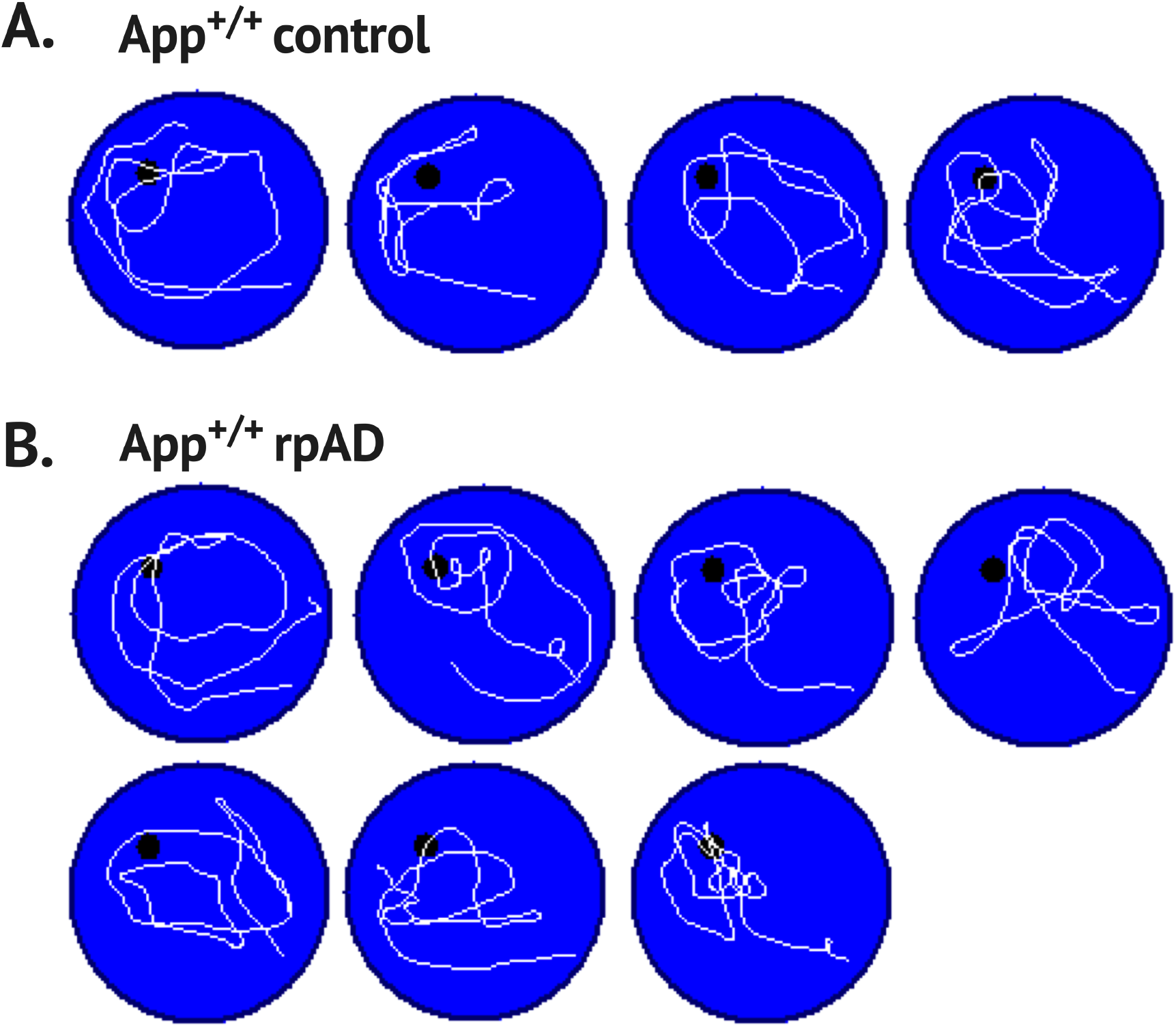
Swim paths for the App^+/+^ control (A) and rpAD (B) seeded mice at three months in the no-platform probe trial

**Fig S1.5.**
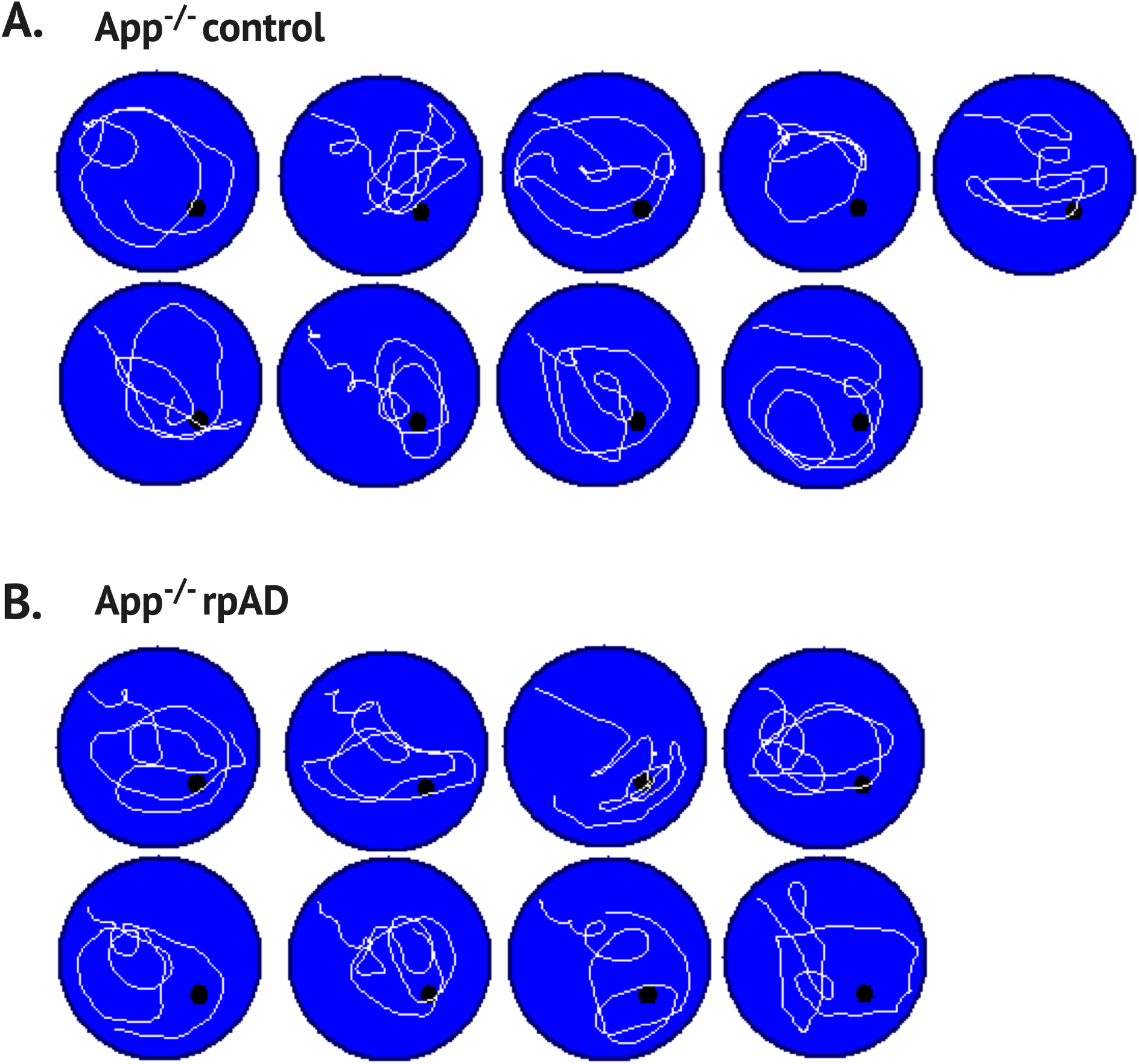
Swim paths for the App^-/-^ control (A) and rpAD (B) seeded mice at six months in the no-platform probe trial

**Fig S1.6.**
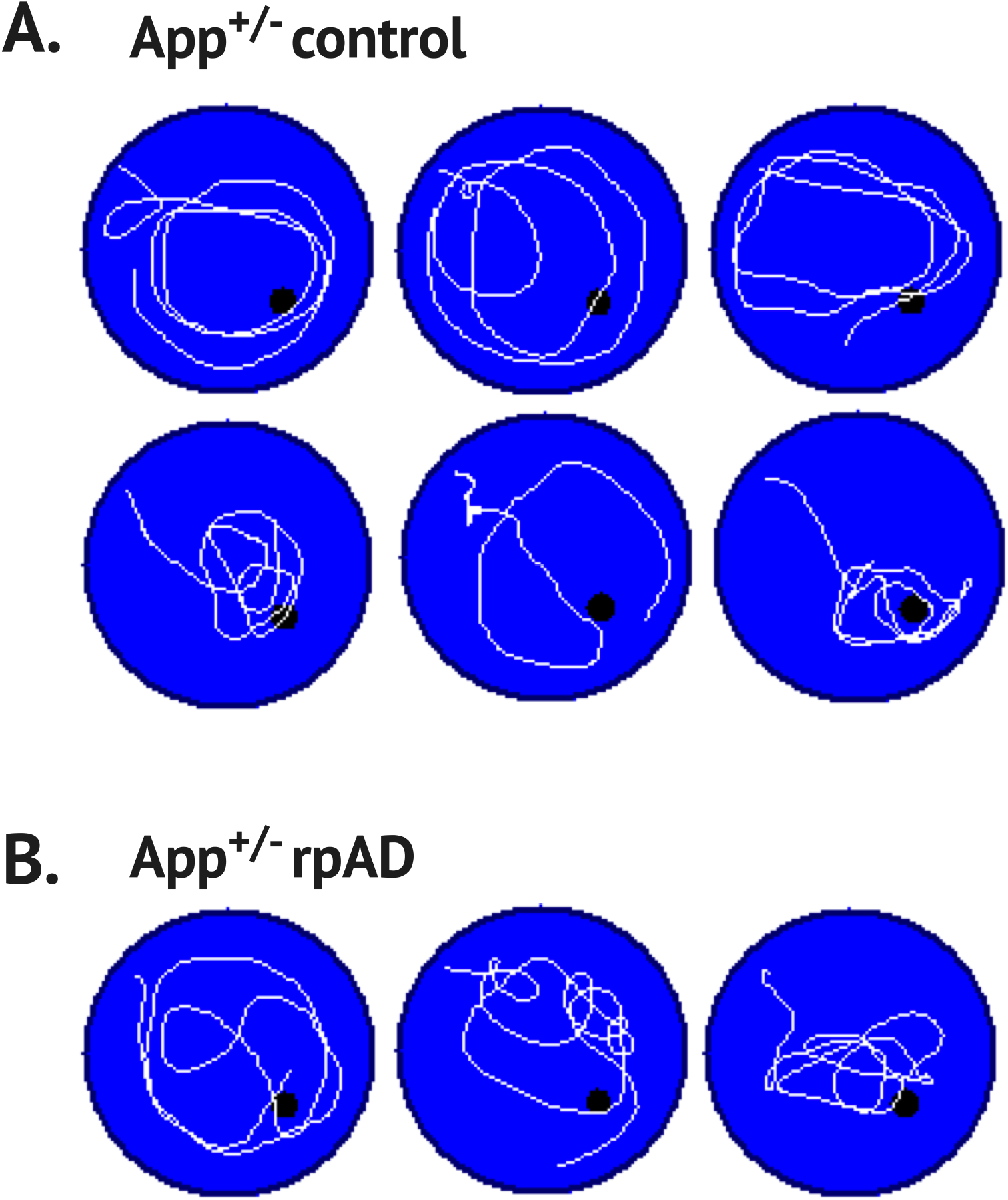
Swim paths for the App^+/-^ control (A) and rpAD (B) seeded mice at six months in the no-platform probe trial

**Fig S1.7.**
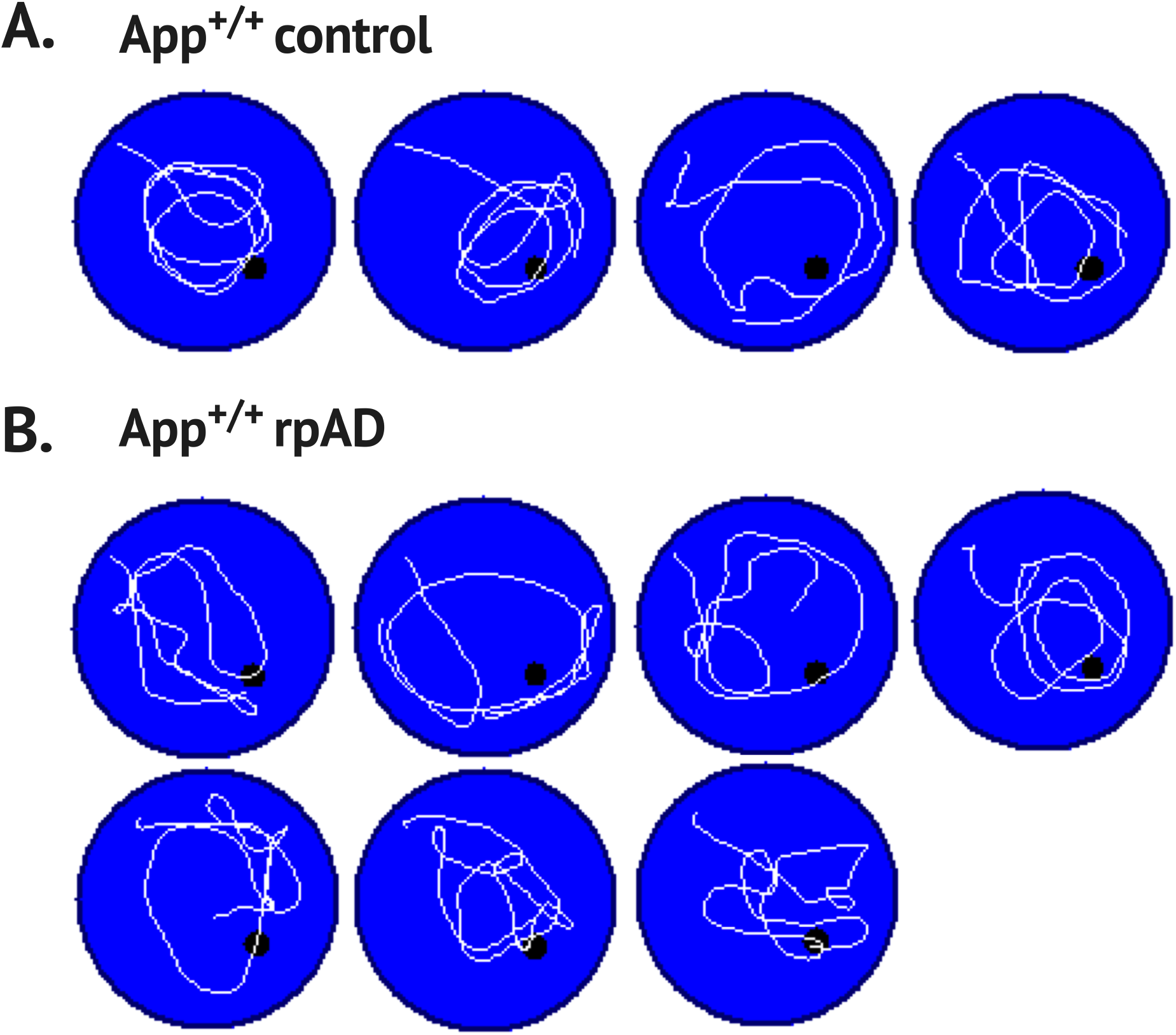
Swim paths for the App^+/+^ control (A) and rpAD (B) seeded mice at six months in the no-platform probe trial

## Notes

### Competing Interest Statement

The authors have declared no competing interest.

